# Diel light cycles affect phytoplankton competition in the global ocean

**DOI:** 10.1101/2021.05.19.444874

**Authors:** Ioannis Tsakalakis, Michael J. Follows, Stephanie Dutkiewicz, Christopher L. Follett, Joseph J. Vallino

## Abstract

Light, essential for photosynthesis, is present in two periodic cycles in nature: seasonal and diel. Although seasonality of light is typically resolved in ocean ecosystem and biogeochemistry models because of its significance for seasonal succession and biogeography of phytoplankton, the diel light cycle is generally not resolved. Here we use a three-dimensional global ocean model and compare high temporal resolution simulations with and without diel light cycles. The model simulates 15 phytoplankton types of different cell size, encompassing two broad ecological strategies: small cells with high nutrient affinity (gleaners) and larger cells with high maximal growth rate (opportunists). Both are grazed by zooplankton and limited by nitrogen, phosphorus and iron. Simulations show that diel cycles of light induce diel cycles in phytoplankton populations and limiting nutrients in the global ocean. Diel nutrient cycles are associated with higher concentration of limiting nutrients by up to 200% at low latitudes (-40° to 40°), a process that increases opportunists’ biomass by up to 50%. Size classes with the highest maximal growth rates from both gleaner and opportunist groups are favored the most by diel light cycles. This mechanism weakens as latitude increases because the effects of the seasonal cycle dominate over those of the diel cycle. The present work shows that resource competition under diel light cycles has a significant impact on phytoplankton biogeography, indicating the necessity of resolving diel processes in global ocean models.

## Introduction

Marine phytoplankton drive photosynthesis and biogeochemical cycles in the ocean, affecting ocean productivity and Earth’s climate. They constitute a polyphyletic group characterized by a tremendous number of species differing in forms and functions (Litchman and Klausmeier, 2008) and spanning more than 7 orders of magnitude in cell volume (Finkel et al., 2010). Although the geographical distribution, or biogeography, of phytoplankton species is still relatively sparsely mapped in the global ocean, major phytoplankton functional groups (e.g. cyanobacteria, diatoms, dinoflagellates) and size classes appear to have distinct differences in biogeography (Barton et al., 2013). Understanding the environmental and ecological factors controlling phytoplankton biogeography is an important element linking community structure with ocean productivity, stability and resilience to environmental stressors (Ptacnik et al., 2008; Winder and Sommer, 2012). Predicting biogeography of phytoplankton size classes is particularly important for climate change predictions, given that large phytoplankton (such as diatoms) have higher sinking rates compared to their smaller counterparts (such as small cyanobacteria), enhancing carbon sequestration from surface to the deep ocean (Tréguer et al., 2018).

To help understand the role of diverse phytoplankton and their biogeography, ocean ecologists have been experimentally derived and constrained parameters that govern the traits and trade-offs of phytoplankton (Litchman and Klausmeier, 2008; Finkel et al., 2010), and these parameters are the foundations for marine ecosystem models (Follows and Dutkiewicz, 2011). Such models have been incorporated into global ocean circulation models to describe general patterns of phytoplankton functional biogeography, including the preponderance of gleaners and opportunists (Aumont et al., 2003; Chai et al., 2007; Dutkiewicz et al., 2009), the realized niche of nitrogen fixers (Dutkiewicz et al., 2014; Landolfi et al., 2015; Follett et al., 2018a) or mixotrophic plankton (Ward and Follows, 2016). Such models have also been used to derive mechanistic explanations on the structure of plankton food webs (Prowe et al., 2012; Ward et al., 2012), biodiversity patterns in the global ocean (Barton et al., 2010; Vallina et al., 2014; Dutkiewicz et al., 2020), as well as the impact of climate change in marine ecosystems (Dutkiewicz et al., 2015; Dutkiewicz et al., 2019).

An important environmental factor considered by the majority of ocean models is the seasonal light cycle (Sommer et al., 2012). The delivery of light and dissolved nutrients to the surface ocean can be highly seasonal and out of phase. At high latitudes, light limitation during winter is associated with low phytoplankton biomass and high residual nutrient concentrations. During spring, availability of both light and nutrients leads to the typical spring bloom period where phytoplankton biomass increases rapidly until nutrients are depleted and limit phytoplankton growth in summer. Although seasonal dynamics and initiation of spring bloom can be controlled by several factors (Behrenfeld and Boss, 2014), the above mechanism describes a major part of phytoplankton seasonal succession, where opportunistic types dominate during the resource-replete spring bloom period and gleaner types prevail in nutrient-depleted conditions in summer. Since seasonality becomes stronger with latitude, ocean models predict an increase in fitness of opportunistic phytoplankton in temperate and polar regions (Follows et al., 2007; Dutkiewicz et al., 2009; Tsakalakis et al., 2018) in agreement with observed patterns in the abundance of diatoms, which are typically considered opportunistic, and pico-cyanobacteria which are small and efficient gleaners (Bracher et al., 2009; Hirata et al., 2011; Acevedo-Trejos et al., 2013; Flombaum et al., 2013).

The diel cycle is the other time scale over which light fluctuates periodically; however, research is limited regarding its effects on phytoplankton biogeography. Resource competition theory suggests that gleaner types should always dominate in a system where resource oscillations are absent or weak (Tilman, 1982; Grover, 1990). Their high nutrient affinity allows them to draw down limiting nutrients to levels where other strategies may not be competitive. However, in the presence of an environment-driven resource oscillation, such as the seasonal cycle, opportunistic types have an advantage during phases of high resource availability based on their higher maximal growth rates (Grover, 1990). Experiments and simple conceptual models have shown an analogous response to diel light cycles and/or light-induced nutrient oscillations, which favor opportunistic phytoplankton over gleaners (Litchman, 1998, 2003; Litchman and Klausmeier, 2001; Litchman et al., 2004). In a previous modelling study we explored the effects of diel light cycles in a simple chemostat system and predicted potential effects on phytoplankton biogeography (Tsakalakis et al., 2018). It was shown that nutrient oscillations driven by diel light cycles favor opportunistic phytoplankton over gleaners in a similar manner to the impact of seasonality, indicating the importance of including both periodic variations of light in marine ecosystem models.

Here we examine the implications of this process in the more complex setting of a global ocean simulation. We ask, to what extent do diel light cycles impact the emergent large-scale biogeography of phytoplankton? We hypothesize that the mechanisms of the idealized study will carry through, but also consider how a more complex physical environment and ecosystem structure affect the impact. We study the effects of diel light cycles in a three-dimensional global ocean circulation and biogeochemical model (Dutkiewicz et al., 2021). The model simulates dynamics of a diverse phytoplankton community with nutrient limitations by nitrogen, phosphorous and iron, where each phytoplankton size class is preyed upon by a dedicated zooplankton consumer. Model results show that diel light cycles induce diel oscillations of limiting nutrients in the global ocean that increase the biomass of fast-growing phytoplankton types, in agreement with the conceptual study of Tsakalakis et al. (2018). Our study indicates that diel light cycles do affect nutrient dynamics and gleaner-opportunist competition and suggests that accounting for diel cycles should improve predictions of marine microbial dynamics and biogeography.

## Methods

The physics of the global ocean model is based on the MITgcm model (Marshall et al., 1997), while the biogeochemical/ecosystem model follows Dutkiewicz et al. (2021) with some modifications that are discussed below. Of significance here, light input is configured to resolve diel cycles and compared to a control simulation without diel light variations.

We use a configuration of the MITgcm ocean model that simulates global ocean circulation, constrained to be consistent with altimetric and hydrographic observations (the ECCO-GODAE state estimates, Wunsch and Heimbach, 2007). It has horizontal resolution of 1° × 1° and 23 vertical levels that span from surface to a maximum depth of 5700 m with level thickness ranging from 10 m near the surface to 500 m at depth. The coupled biogeochemical/ecosystem model resolves the cycling of carbon, nitrogen, phosphorus, iron and oxygen through inorganic, living, particulate and dissolved organic constituents. It resolves 15 phytoplankton types differing in size and biogeochemical function and 15 size classes of zooplankton. Phytoplankton types are spaced uniformly in log space from 0.6 to 228 μm equivalent spherical diameter (ESD) and grouped into two functional groups: the 4 smallest are gleaners (0.6 to 2.0 μm ESD), while the remaining 11 are opportunists (3.0 to 155 ESD). For ease in interpretation, our model community is simplified compared to Dutkiewicz (2021); the gleaners are analogs of pico-cyanobacteria and pico-eukaryotes, while the opportunists are based on diatom growth parameters, except they are not limited by silica here.

Phytoplankton parameters such as maximum growth rate, affinity for nutrient uptake, grazing by zooplankton and sinking are parameterized as a power-law function of cell volume informed by compilations of phytoplankton growth parameters (Ward et al., 2012; Dutkiewicz et al., 2020). Phytoplankton maximum specific growth rates have a unimodal relationship with cell size peaking at intermediate sizes, in agreement with experimental observations (Raven, 1994; Finkel et al., 2010; Marañón et al., 2013; Sommer et al., 2017; Dutkiewicz et al., 2020); phytoplankton smaller than 3 μm have an increase in specific growth rate with size while those larger than 3 μm have a decrease in specific growth rate with size. It is also assumed that affinity for nutrient uptake decreases with cell size; the smallest phytoplankton have the highest affinity for nutrient uptake (Edwards et al., 2012) based on their higher surface-to-volume ratio (Kiørboe, 1993; Raven, 1994).

The model uses Monod kinetics to describe nutrient dependent growth of phytoplankton (Monod, 1949), while C:N:P:Fe requirements are constant but different among functional groups. Chl *a* for each phytoplankton type is variable and dependent on light and nutrient availability, following Geider et al. (1998). Zooplankton grazing is resolved using a Holling type III grazing function (Holling, 1959) for 15 zooplankton size classes (from 6.6 to 1635 μm ESD) that graze on phytoplankton 10 times smaller than themselves. Maximum grazing rate decreases with size following Hansen et al. (1997). In Dutkiewicz et al. (2021), each zooplankton size class can prey upon several phytoplankton size classes and zooplankton smaller than themselves. Here we simplify predation so that a given zooplankton size class grazes on a single phytoplankton type and does not graze upon other zooplankton. This simplification allows for a better interpretation of the bottom-up effects driven by diel light cycles, avoiding complex trophic interactions and cascading effects associated with more sophisticated grazing configurations.

Two light regimes were considered, a control simulation with only seasonal light cycles and a diel simulation with both seasonal and diel light cycles. In the control, monthly averaged irradiance levels in the surface ocean were provided by the Ocean-Atmosphere Spectral Irradiance Model (OASIM) that includes the impact of clouds, water vapor and aerosols in the atmosphere, as used in Dutkiewicz et al. (2020), but here we sum the different light wavelengths providing a single dimension of photosynthetically active radiation (PAR). In the diel simulation, Brock’s (1981) system of equations was used to produce surface light fields with high temporal resolution. Brock’s equations calculate diel cycles of light intensity and changes in day length across the globe’s surface, unperturbed by cloud formation or other processes. The light field with diel cycles was then normalized to match the annual average irradiance of the control simulation. This ensured that the two simulations have the same light energy input on average, allowing for a fair comparison between them. Model runs of 10 years with a time step of 3 h^-1^ were used, while test runs of 20 years ensured that the 10-year runs are sufficient for reaching a quasi-steady state in terms of plankton biogeography, which was typically achieved between the first 2 to 5 model years. The last simulated year is used to analyze annually averaged patterns and annual and diel dynamics of model variables.

When using the same growth parameter values between control and diel simulations, the diel case leads to substantially lower primary production; the average of a non-linear function (the diel simulation in this case) is not usually equal to the non-linear function with averaged input (the control simulation). This is known as Jensen’s inequality (Jensen, 1906; Denny, 2017). Particularly to our model analysis, nighttime increases phytoplankton mortality in the diel simulation while the higher light intensity during the day (compared to the control simulation) cannot compensate night loses. In order to match primary production of the diel simulation to the control, we increased maximal specific growth rates of all phytoplankton types by a factor of 2.05 (we tried several factors and chose the factor that matches the control simulation, see Fig. 1). With this alteration other model variables, such as limiting nutrients and plankton biomass, also approach concentrations found in the control simulation (Fig. S1). The increase in specific growth rate for the diel case allows for a fair comparison between diel and control simulations. Even though the maximal growth rates of phytoplankton types are different between simulations, we ensure that nutrient affinity for phytoplankton growth kinetics remains constant using the following equation (see Follows et al., 2018):

**Figure 1.**
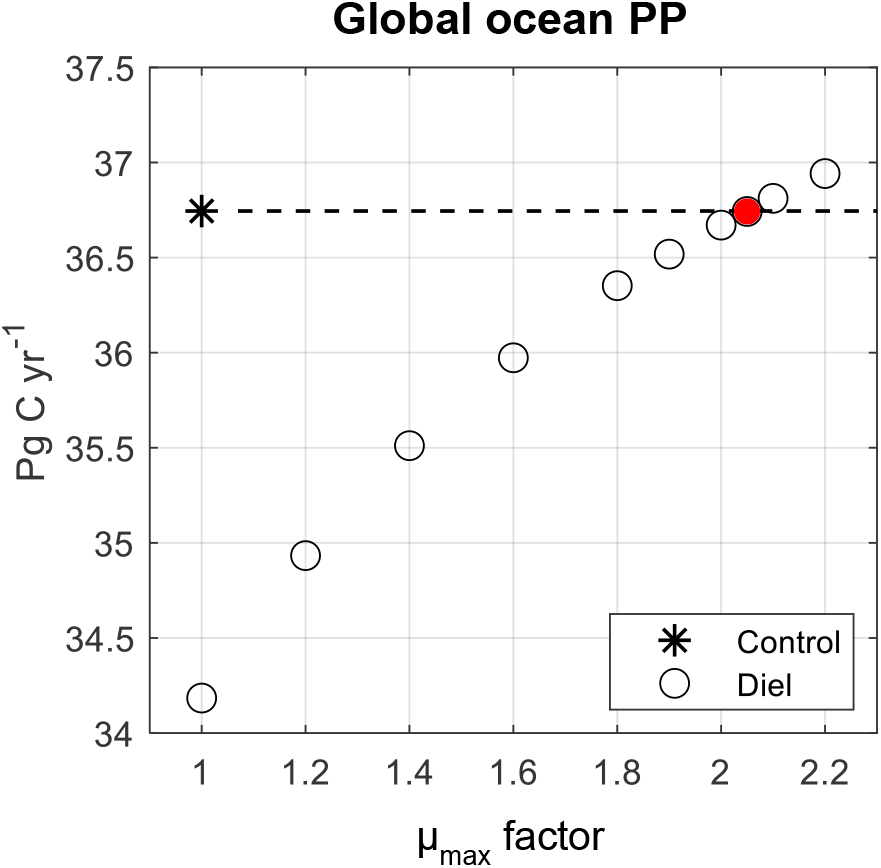
Global ocean primary production (PP) of model simulations with only seasonal light cycles (Control) and both seasonal and diel light cycles (Diel). The 10 diel simulations differ in phytoplankton growth parameterization; maximal growth rates of all phytoplankton types are multiplied by an μ_max_ factor indicated in the x-axis. Note that the diel simulation with μ_max_ factor = 1 is parameterized with same maximal growth rates as the control. The dashed line is an extension of the control, while the red circle highlights the μ_max_ factor = 2.05 where the diel simulation matches the control in global ocean PP.

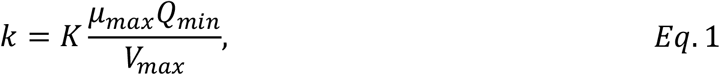

where *K* is the half saturation constant for growth used in the Monod growth model, *K* is the half saturation constant for nutrient uptake, *μ*_*max*_ the maximal growth rate, *Q*_*min*_ the minimum nutrient quota for growth, and *V*_*max*_ the maximum nutrient uptake rate. According to the equation above, half saturation for growth is proportional to the maximal growth rate so that growth affinity 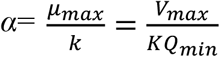 is independent of *μ*_*max*_.

To illustrate the effect of diel light cycles on a model variable *X*, such as the concentration of a limiting nutrient or a phytoplankton type/group, we use the following metric:

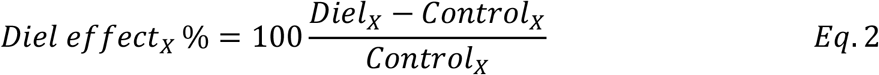

where *Diel*_*x*_ and *Control*_*x*_ are the concentrations of variable *X* in diel and control simulations, respectively.

## Results

The model captures the patterns of high productivity in the equatorial and subpolar regions, and the oligotrophic conditions in subtropical gyres (Fig S2, model results can be retrieved online, Tsakalakis et al., 2021). The control simulation shows typical distributions of limiting nutrients in the global ocean (Fig. 2a to c). Dissolved inorganic nitrogen (DIN = NO_3_^-^+NO_2_^-^+NH_4_^+^) has low concentrations in the Indian and Atlantic Oceans as well as the subtropics of the Pacific Ocean (Fig. 2a). Low concentrations of phosphate follow a similar pattern to DIN (Fig. 2b), while both DIN and PO_4_^3-^ are in excess in the Southern Ocean, as this area is known to be primarily limited by iron and light (Thomas, 2003). Iron exhibits low concentrations in the Pacific and Southern Oceans, while it reaches high concentrations in the Indian and Atlantic Oceans (Fig. 2c).

**Figure 2.**
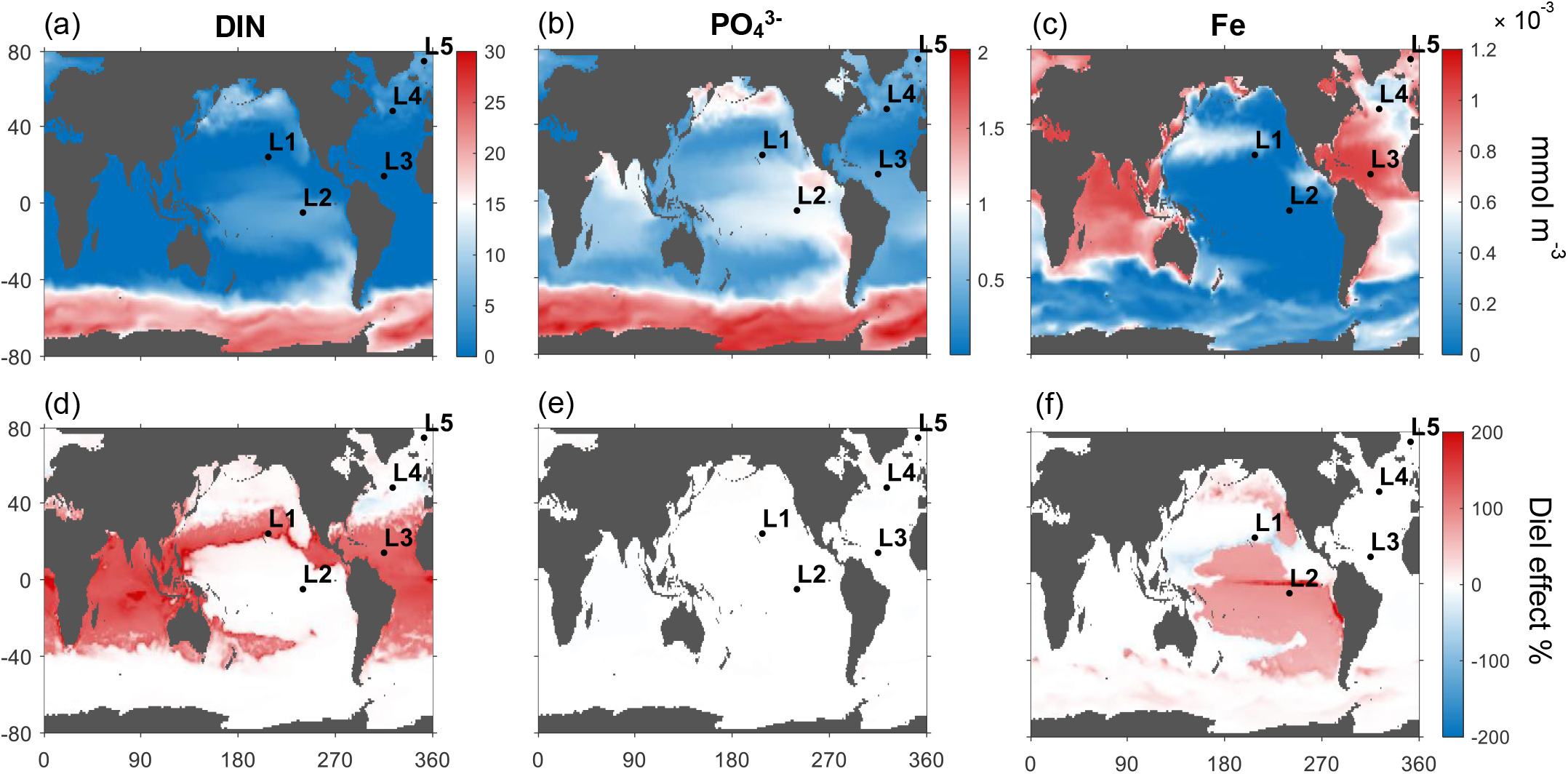
Effect of diel light cycles on concentrations of limiting nutrients. The upper panel shows annually averaged surface (0 – 55 m depth) nutrient concentrations in the control simulation and the lower panel shows the effect of diel light cycles compared to the control (Eq. 2, difference in concentration between diel and control simulations, divided by the control, %). DIN refers to the sum of NO_3_^-^, NO_2_^-^ and NH_4_+. Labelled black dots indicate locations (L1 to L5) for which annual dynamics are presented in Fig.3.

Diel light cycles increase the concentration of the most limiting nutrient at low latitudes. We focus on model results of the surface ocean (0 to 55 m depth) because the effects driven by diel light cycles are restricted to the surface (Fig. S2). In Fig. 2d to f, we present the effect of diel light cycles on nutrient concentrations compared to the control (Eq. 2). Overall, increases of nutrient concentrations are mainly restricted to low latitudes (-40° to 40°). Diel light cycles increase DIN concentrations up to 200% in regions where DIN has its lowest concentration in the control, i.e., low latitudes in Indian and Atlantic oceans and subtropics of the Pacific (Fig. 2d). Fe concentrations increase significantly (up to 200% as well) in the Pacific Ocean, especially at low latitudes (Fig. 2f). Note that the patterns of increased concentrations of DIN and Fe are complementary, i.e., only a single nutrient increases locally. Calculation of nutrient limitation for gleaner and opportunist types shows that the distribution of DIN and Fe limitation coincides with the increases of those nutrients in the diel simulation, supporting that diel light cycles increase the local concentration of the most limiting nutrient (Fig. S3). In contrast, PO_4_^3-^ concentrations are not affected by diel light cycles (Fig. 2e), because PO_4_^3-^ is not the most limiting nutrient at any low-latitude region in the model (Fig. S3).

We present model timeseries in 5 ocean locations (Fig. 3) ranging from low latitudes (L1 to L3, locations shown in Fig. 2) to a temperate (L4) and a polar location (L5). These five locations summarize the general effects of adding diel light cycles. Dynamics of additional locations distributed across the global map are presented in Fig. S5. Photosynthetically active radiation (PAR) exhibits (as per experiment design) only seasonal cycles in the control simulation (red lines, Fig. 3, row 1) and both seasonal and diel cycles in the diel simulation (grey lines, Fig. 3, row 1).

**Figure 3.**
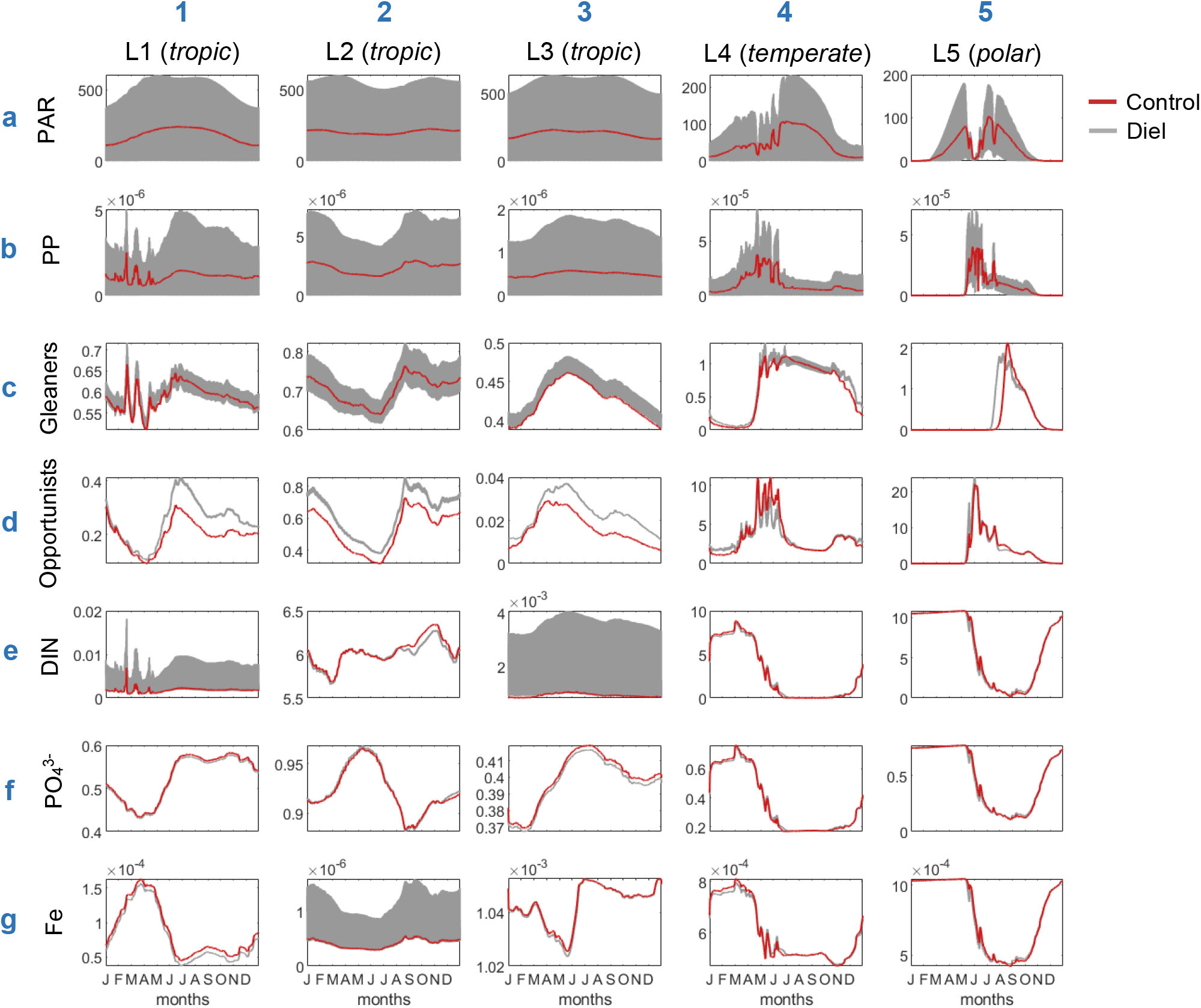
Annual dynamics in the control (red) and diel (grey) simulations of photosynthetically active radiation, PAR (μEinstein m^-2^ s^-1^), primary production, PP (mmol C m^-3^ s^-1^), biomass of phytoplankton gleaners and opportunists (mmol C m^-3^), DIN = NO_3_^-^, NO_2_^-^ and NH_4_+ (mmol N m^-3^), PO_4_^3-^ (mmol P m^-3^) and Fe (mmol Fe m^-3^) at 5 locations of the surface ocean (averaged from 0 to 55 m depth). Locations (L1 to L5) are illustrated in Fig. 2. Note that grey shaded areas are due to diel oscillations.

Dynamics at low latitudes (L1 to L3) show that diel light cycles induce strong oscillations and higher concentrations of the most limiting nutrient (DIN in L1 and L3, and Fe in L2). PAR diel cycles at the low-latitude locations (Fig. 3 a1 to a3) induce pronounced diel oscillations throughout the year in primary production (Fig. 3 b1 to b3) and in the biomass of gleaners and opportunists (Fig. 3 c1 to c3 and d1 to d3, respectively). Those diel cycles of limiting nutrients in tropics result in higher biomass for opportunists, while the averaged biomass of gleaners is not significantly affected despite its strong diel oscillations.

Dynamics at higher latitudes (locations L4, temperate and L5, polar) show that the impact of seasonality dominates over the effects of diel light cycles; nutrient and biomass levels are very similar between control and diel simulations (Fig 3, column 4 and 5), oscillating strongly at a seasonal scale. Although diel nutrient cycles are still present at temperate and polar locations in the diel simulation, they are barely discernable in the figures as seasonal oscillations are much larger in amplitude. Several locations at temperate latitudes across the global map support the same result (Fig. S5).

The diel cycle in limiting nutrients is a result of cessation of photosynthesis (and nutrient uptake in our Monod-based model) during the night, but the continuation of remineralization of detrital matter (Fig 4c). During daytime in the diel simulation, phytoplankton nutrient uptake decreases nutrient levels towards the concentration of the control simulation. This mechanism of accumulation of nutrients at night time maintains the higher concentrations (up to 200%) in DIN and Fe than in the control (Fig. 2d and f). PAR and PP also have a strong diel pattern in the diel simulation, as expected (Fig. 4a and b). Note that PAR slightly decreases during the day based on the increasing phytoplankton biomass that absorbs light (grey line, Fig. 4a), while PP also declines during daytime based on the gradual increase in nutrient limitation (grey line, Fig. 4b).

**Figure 4.**
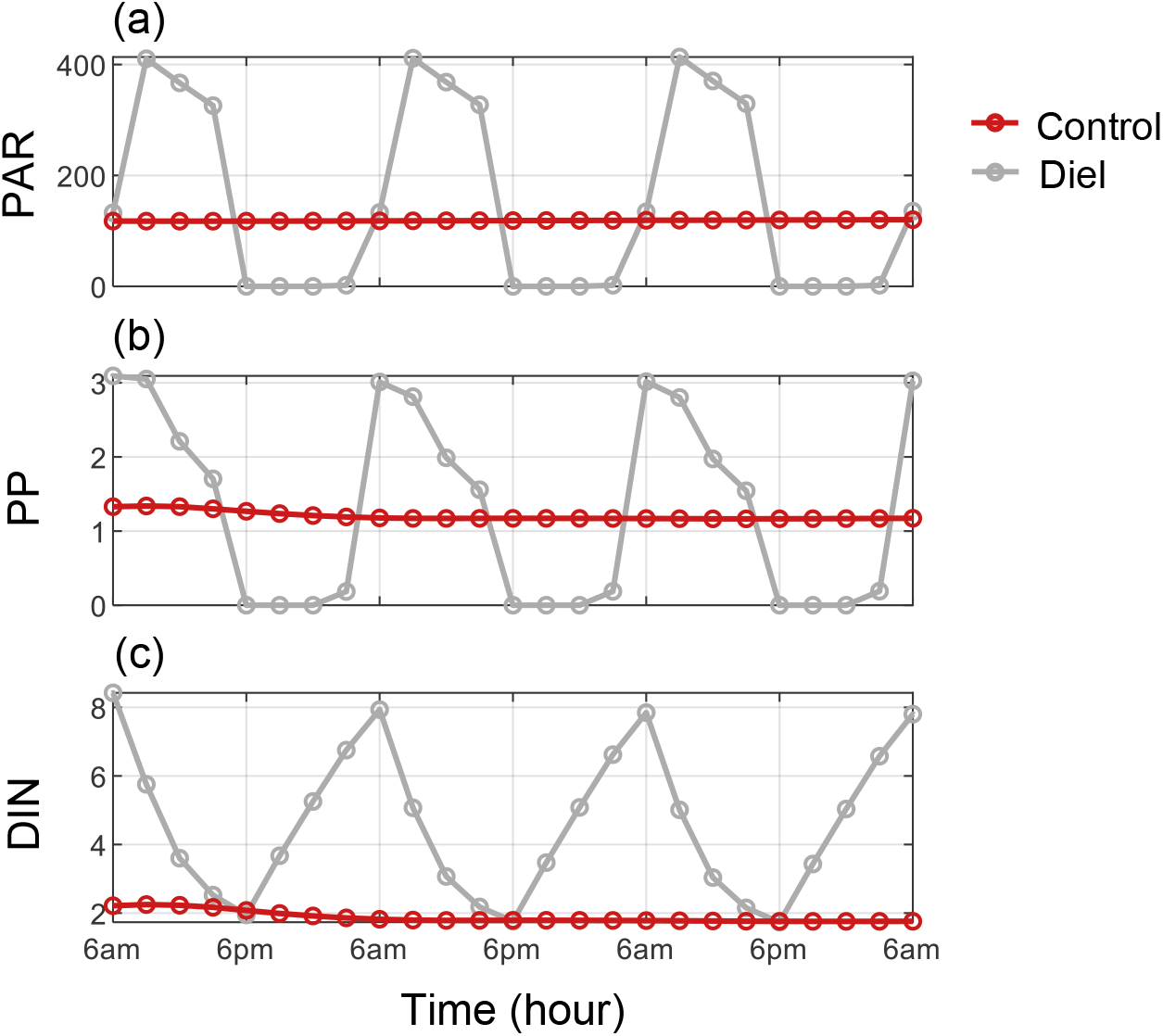
Diel dynamics in the control (red) and diel (grey) simulations of the surface ocean (0 to 55 m depth) for three days, from January 22 to 24, at the tropical location L1 (Fig. 2): (a) Photosynthetically active radiation, PAR (μEinstein m^-2^ s^-1^); (b) primary production, PP (nmol C m^-3^ s^-1^); (c) DIN = NO_3_^-^, NO_2_^-^ and NH_4_+ (μmol N m^-3^). Circles indicate model time step equal to 3h^-1^.

Opportunists are favored by the presence of diel nutrient cycles at low latitudes (Fig. 5). The biogeography of gleaners and opportunists in the control simulation follows a similar pattern as previous studies, with biomass of gleaners more uniformly distributed across the global ocean (Fig. 5a) while opportunists have low concentrations at low latitudes, but substantially increased biomass in temperate and subpolar regions (Fig. 5b). Note that areas with ice presence are masked out (white with hatching), as phytoplankton biomass underneath ice is generally at very low concentrations and sensitive to our diagnostics. Addition of diel light cycles in the model increases opportunists’ biomass by up to 50% compared to the control (Fig. 5d), a pattern that is restricted to low latitudes where significant diel cycles in limiting nutrients are present. By contrast, gleaners do not show any significant change at low latitudes (Fig. 5c); opportunistic species with higher maximal growth rates are better adapted to utilizing the temporary high nutrient concentrations. The timeseries results (Fig. 3) in the tropical locations L1 to L3 show that diel light cycles affect phytoplankton competition during all seasons; higher nutrient concentrations due to diel light cycles result in higher biomass of opportunists throughout the year.

**Figure 5.**
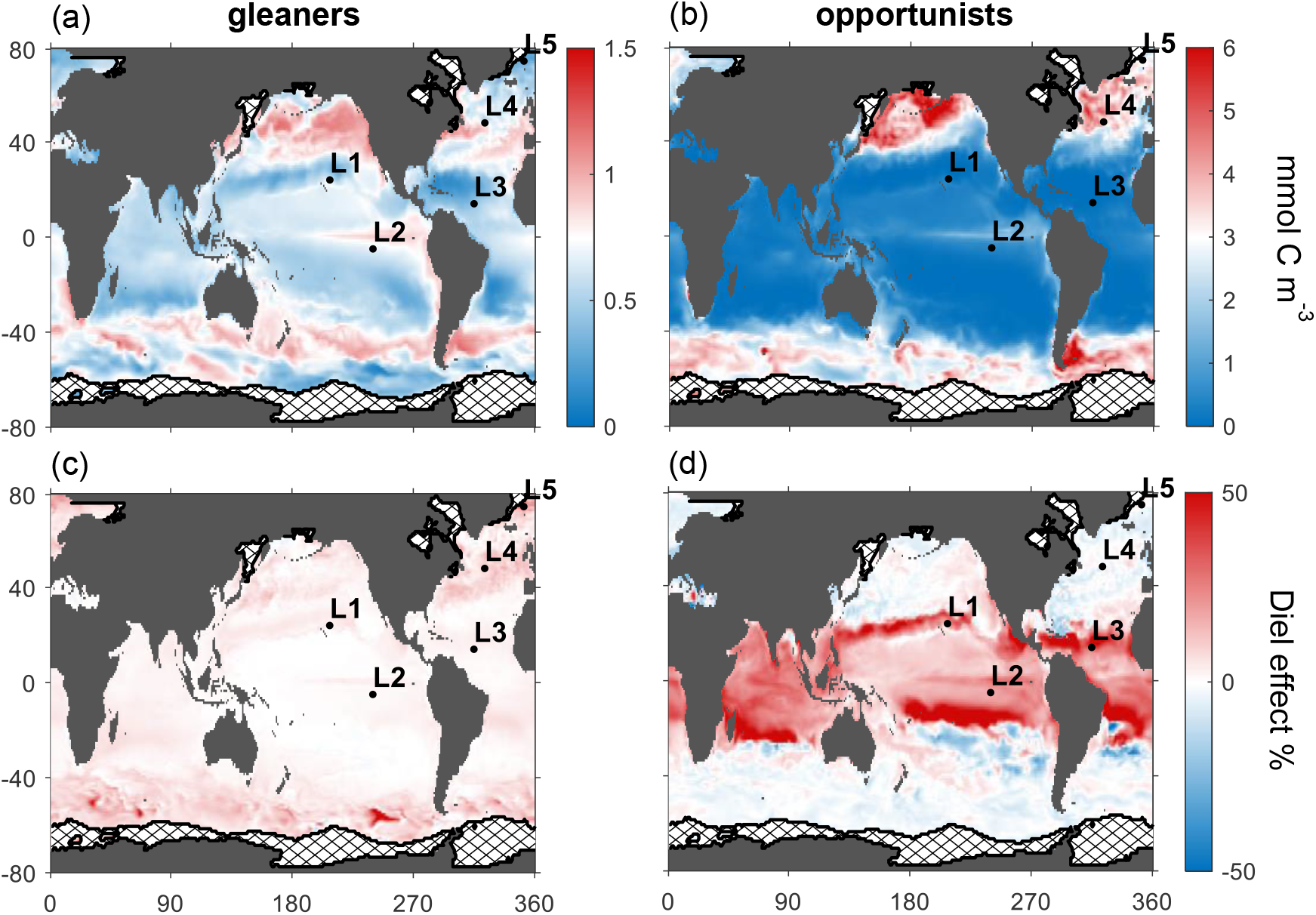
Effect of diel light cycles on phytoplankton biogeography. The top row shows annually averaged surface concentrations (0 – 55 m depth) of gleaners and opportunists in the control simulation and the bottom row shows the effect of diel light cycles compared to the control (Eq. 2, difference in biomass between diel and control simulations, divided by the control, %). Labelled black dots indicate locations (L1 to L5) of which dynamics are presented in Fig. 3. White with hatching refers to areas with ice presence.

Analysis of biomass changes associated with diel light cycles of the 15 phytoplankton types indicates that fast-growing size classes are favored at low latitudes (Fig 6a and b). Note that the largest gleaners and the smallest opportunists have the highest maximal growth rates among their groups. In tropical regions of both South and North hemispheres, the smallest opportunists increase in biomass in the diel simulation compared to the control (red circles, Fig. 6a and b). This pattern persists but becomes weaker at temperate latitudes (Fig. 6c and d). The largest gleaners, that have the highest maximum specific growth rate among gleaners, also slightly increase in tropical regions in the diel simulation (Fig. 6a and b). At temperate latitudes, all gleaner sizes increase in biomass (Fig. 6c and d); this result cannot be understood by nutrient dynamics, but it is rather an artifact of our model parameterization discussed in the paragraph below. Overall, Fig. 6 illustrates that diel light cycles favor fast-growing types from both gleaner and opportunist groups at low latitudes, where diel cycles in limiting nutrients are present.

**Figure 6.**
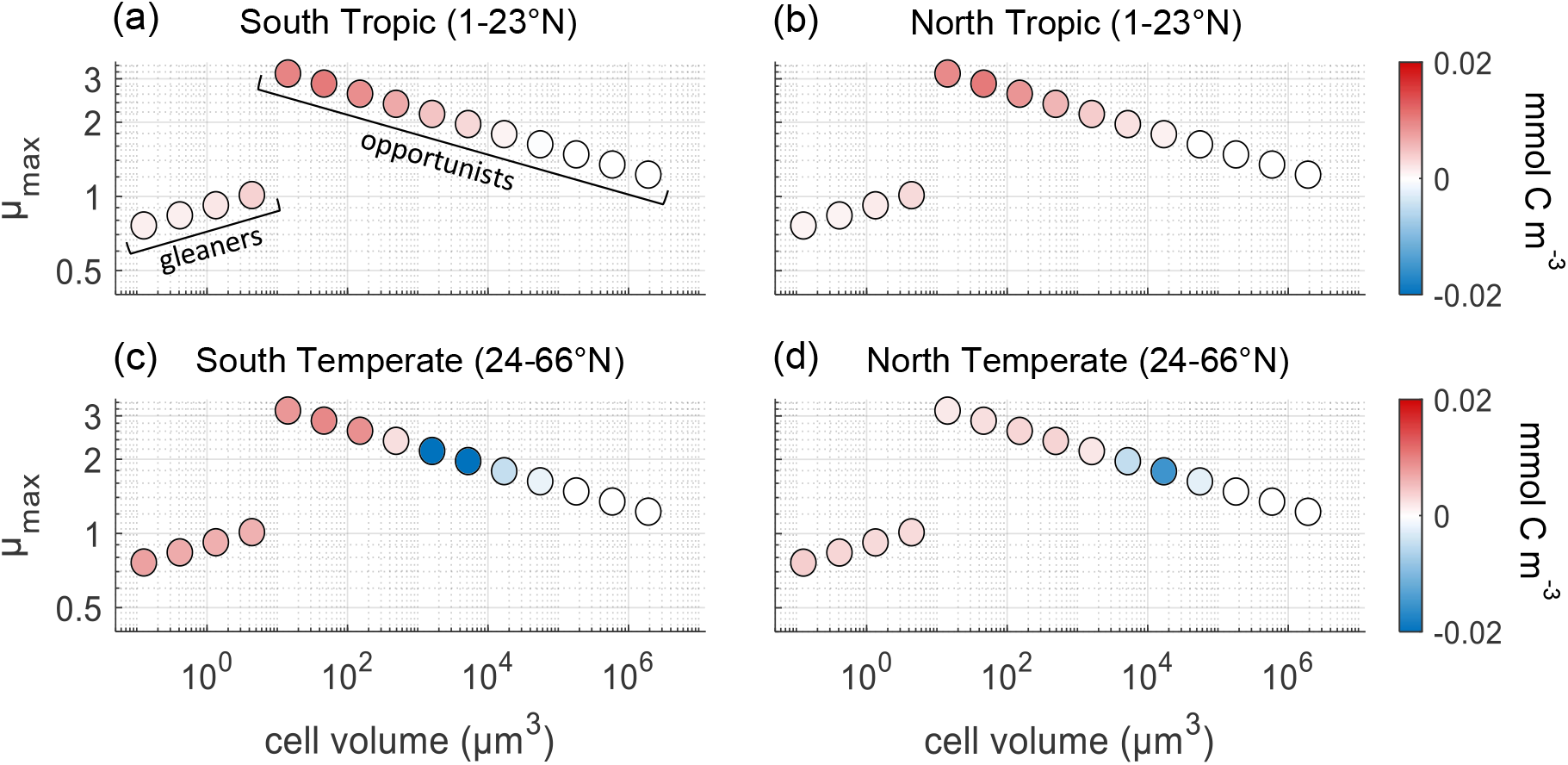
Effect of diel light cycles on phytoplankton size classes in tropical and temperate regions. The 15 modelled phytoplankton types are shown (circles) as a function of their maximal growth rate of the control simulation (μ_max_, d^-1^) versus cell volume. Coloring refers to the average concentration (0 – 55 m) difference between simulations (diel – control) for each phytoplankton type in the respective latitudinal region. Phytoplankton groups of gleaners and opportunists are annotated in the top-left diagram.

Phytoplankton gleaners perform slightly better than opportunists at high latitudes when diel light cycles are added (Fig. 5c and d). This pattern cannot be supported by a resource-competition mechanism, given that nutrient dynamics in temperate and polar regions are very similar between the control and diel simulations (Fig. 3, column 4 and 5). We find that by increasing the maximal growth rate of all phytoplankton to match the global primary production of the control (Fig. 1), we slightly favor the performance of gleaners over opportunists. This is because gleaners are parameterized with low maximal growth rates compared to opportunists, so their relative fitness is better at higher μ_max_ factors. This is evident in the biogeography plots (Fig. 5c and d), in which the global response to diel cycles of gleaners is always positive but opportunists are both positively and negatively affected.

## Discussion

Global ocean models aim at capturing large scale patterns of community structure and ocean biogeochemistry. To achieve that, the essential environmental factors and ecological interactions should be considered. The goal of the present study is to demonstrate the effects of the diel light cycle on phytoplankton competition in the global ocean. Based on the theory of resource competition in oscillating environments developed in the chemostat system, we expect that diel light cycles induce cycles in limiting nutrients in the ocean, a process that favors fast-growing phytoplankton (Litchman and Klausmeier, 2001; Tsakalakis et al., 2018). Here we show that this mechanism persists in a global ocean model which resolves several realistic features of marine ecosystems, such as resource competition in the water column, multi-nutrient limitation, spatial and temporal variation of resource supply and zooplankton predation.

Diel nutrient cycles in the surface ocean are caused by increases during the night due to remineralization and decreases in the day due to uptake (Fig. 3, 4). This mechanism significantly increases the concentrations of limiting nutrients (up to 200%) at low latitudes (Fig. 2) that favors opportunistic phytoplankton over gleaners (Fig. 5). In the control simulation, limiting nutrients are maintained at low concentrations at these low latitudes (Fig. 3) because the weak seasonality allows phytoplankton to deplete limiting nutrients throughout the year. Under such conditions, gleaner types dominate due to their high nutrient affinity compared to other strategies (lowest R*, following terminology of Tilman, 1982), but this competitive advantage weakens under diel nutrient fluctuations. (Note though that opportunists do not necessarily become dominant). Non-limiting nutrients also oscillate with a diel frequency in the model, however, they are not maintained at low concentrations by phytoplankton. Therefore, diel oscillations of non-limiting nutrients are not significant relative to other sources of variability such as seasonal dynamics (even in the weak seasonality of tropics) and spatial nutrient circulation. This pattern of limiting and non-limiting nutrient dynamics is consistent for several locations across the global ocean (Fig. 3, S5).

The amplitude of diel nutrient oscillations in our model is up to 0.1 mmol N m^-3^ for DIN, 8 ∗ 10^−3^ mmol P m^-3^ for phosphate and 5 ∗ 10^−6^ mmol Fe m^-3^ for iron. Those oscillations seem small in comparison to seasonal nutrient variations; however, our model analysis shows that they significantly affect phytoplankton competition at low latitudes where limiting nutrients are at low concentrations throughout the year. It is important to validate modelled diel nutrient cycles against ocean observations, but such data are very sparse. The lack of data is because concentrations of limiting nutrients are often at undetectable levels, especially at low-latitude regions where the diel effect is more important. However, diel cycles in nitrate concentrations have been captured using automated measuring methods in coastal ecosystems (Johnson et al., 2006; Sakamoto et al., 2017) and the open ocean (Zhang et al., 2001; Johnson et al., 2013). The study of Zhang et al. (2001) in a marine anticyclonic eddy in the North Atlantic shows that surface nitrate oscillates with an amplitude of 0.045 mmol N m^-3^ peaking at night and declining during the day, supporting the mechanism proposed in our study (Fig. 4). The study area of Zhang et al. (2001) is in the same region as location L4 of our analysis, where our model presents nitrate oscillations with an amplitude of 0.025 mmol N m^-3^ during the same time-period (Fig. 3 e4), showing that our model underestimates the amplitude in that case. This indicates that the effect of diel cycles on phytoplankton competition might be even stronger than our current model predictions, but more observations at several ocean locations are needed for model validation.

Our model also predicts diel cycles in phytoplankton populations, especially at low latitudes where seasonal dynamics are weak (Fig. 3). Observations confirm that the biomass of marine phytoplankton oscillates with a diel pattern (Fuhrman et al., 1985; Stramska and Dickey, 1992; Boysen et al., 2020). The study of Boysen et al. (2020) captures diel cycles in pico-phytoplankton populations (*Prochlorococcus sp*., *Synechococcus sp*. and pico-eukaryotes) near Hawaii. Note that location L1 in our model is also near Hawaii (Fig. 3), and the four smallest phytoplankton types in our model (gleaners) correspond to a *Prochlorococcus* type, a *Synechococcus* type and two types representing pico-eukaryotes. Boysen et al. (2020) report diel population cycles with an amplitude up to ∼0.125 mmol C m^-3^ for *Prochlorococcus sp*., *∼*0.008 mmol C m^-3^ for *Synechococcus sp*. and ∼0.08 mmol C m^-3^ for pico-eukaryotes, while for the same functional groups our model predicts amplitudes of 0.13, 0.1 and 0.1 mmol C m^-3^, respectively. Although our model overestimates the amplitude of diel cycles in *Synechococcus sp*., it does capture the amplitude of the diel population cycles for *Prochlorococcus sp*. and pico-eukaryotes.

The model suggests that diel light cycles significantly affect phytoplankton biogeography by increasing the biomass of opportunistic phytoplankton at low latitudes (Fig. 5d); the smallest size classes of opportunists (range of nano-phytoplankton) are favored the most, as they have the highest maximum specific growth rates among size classes (Fig. 6). Phytoplankton competition at low-latitude oceans inferred by remote-sensing observations indicates that pico-phytoplankton (such as *Prochlorococcus sp*.) dominate at subtropical oligotrophic gyres, while nano-phytoplankton (such as diatoms) dominate at more productive tropical waters (Bracher et al., 2009; Brewin et al., 2010; Hirata et al., 2011). Previous modelling studies have shown that resistance to predation might be a trait explaining the presence, but not dominance, of nano-phytoplankton at low latitudes (Prowe et al., 2012; Ward et al., 2012). Our study highlights an additional and potential bottom-up mechanism taking place at low latitudes that increases the competitive ability of opportunists, or nano size classes. This mechanism might be significant for climate change predictions given that larger phytoplankton, such as diatoms, enhance export of carbon from the surface to the deep ocean (Tréguer et al., 2018).

Furthermore, the model shows that diel light cycles may affect the differences in biogeography among pico-phytoplankton populations. The largest gleaners, which have the highest maximum specific growth rate among members of the gleaner’s group in our model, increase their abundance at low latitudes when diel light cycles are added in the model (Fig. 6a and b). This information is relevant for biogeography studies of keystone pico-phytoplankton populations, such as *Prochlorococcus sp*. versus *Synechococcus sp*. (Flombaum et al., 2013), or ecotypes of *Prochlorococcus sp*. (Martiny et al., 2009). Our work indicates that phytoplankton gleaners with higher maximum specific growth rates might be favored by diel nutrient cycles at low-latitude oceans.

### Future perspectives

The best approach to estimate the impact of diel light cycles in the global ocean would be to develop a model with experimentally derived growth parameters of phytoplankton under diel oscillating conditions. Although there are meta-analyses studies determining growth parameters of several phytoplankton (Edwards et al., 2012, 2015; Finkel et al., 2010), maximal growth rates are typically estimated per day, thus, ignoring the impact of diel cycles. Our analysis shows that when accounting for diel light cycles, the maximal growth rate of phytoplankton should be roughly doubled to produce levels of ocean primary production found using daily averaged growth rate (Fig. 1). This is understandable, as diel light cycles require phytoplankton to photosynthesize only during the daytime, which is roughly half of the day compared to conditions of averaged diel light input that is used in most ocean models. Our work indicates that laboratory studies that characterize the diel cycle of growth rates would be helpful for future ocean models that resolve diel processes.

A relevant extension of the proposed model is to include nutrient storage by phytoplankton with the use of a cell-quota model (e.g. Ward et al., 2012). In the Monod model used here, photosynthesis and nutrient uptake are coupled; carbon and nutrients are instantly converted into biomass when they are both taken up by phytoplankton. Therefore, in a model resolving diel light cycles, uptake of nutrients is only possible during the day. Nutrient uptake by phytoplankton is often found significantly higher during the day than at night (Cochlan et al., 1991; Glibert and Garside, 1992; Yingling et al., 2021) so that a Monod model seems relevant to capture the general diel dynamics. However, studies have shown that some phytoplankton continue to uptake nutrients during nighttime and store them in internal nutrient pools (Needoba and Harrison, 2004; Follett et al., 2018b), while large phytoplankton have significantly larger internal nutrient pools than smaller cells (Marañón et al., 2013). This indicates that the diel cycle may help explain the presence of large phytoplankton specializing in storing nutrients during nighttime.

The circadian clock is also a well-known trait improving microbes’ fitness under diel light cycles (Johnson et al., 2017; Kolody et al., 2019), which can be either strongly present in phytoplankton species (Cohen and Golden, 2015) or absent in others (Holtzendorff et al., 2008). This suggests that some phytoplankton may strongly depend on the diel cycle for their survival or competitiveness, while others do not. A recent work shows that the circadian clock as well as nutrient storage both increase energy acquisition by phytoplankton growing in a chemostat model under diel light cycles (Vallino and Tsakalakis, 2020), supporting the potential of those traits to affect phytoplankton competition and biogeography in the global ocean.

## Acknowledgements

We would like to thank Oliver Jahn for his help on model simulations. Simons Collaboration on Computational Biogeochemical Modeling of Marine EcosystemS supported MJF and SD on CBIOMES grant #549931; CLF on CBIOMES grants #827829 and #553242; and JJV and IT on CBIOMES grant #549941. The National Science Foundation supported IT and JJV on award OCE-1558710 and JJV on awards OCE-1637630 and DEB-1655552.

## Data availability statement

The data that support the findings of this study are openly available in ‘Global ocean model results – effects of diel light cycles’ at http://dx.doi.org/10.5281/zenodo.4772367.

## Supplementary figures

**Figure S1.**
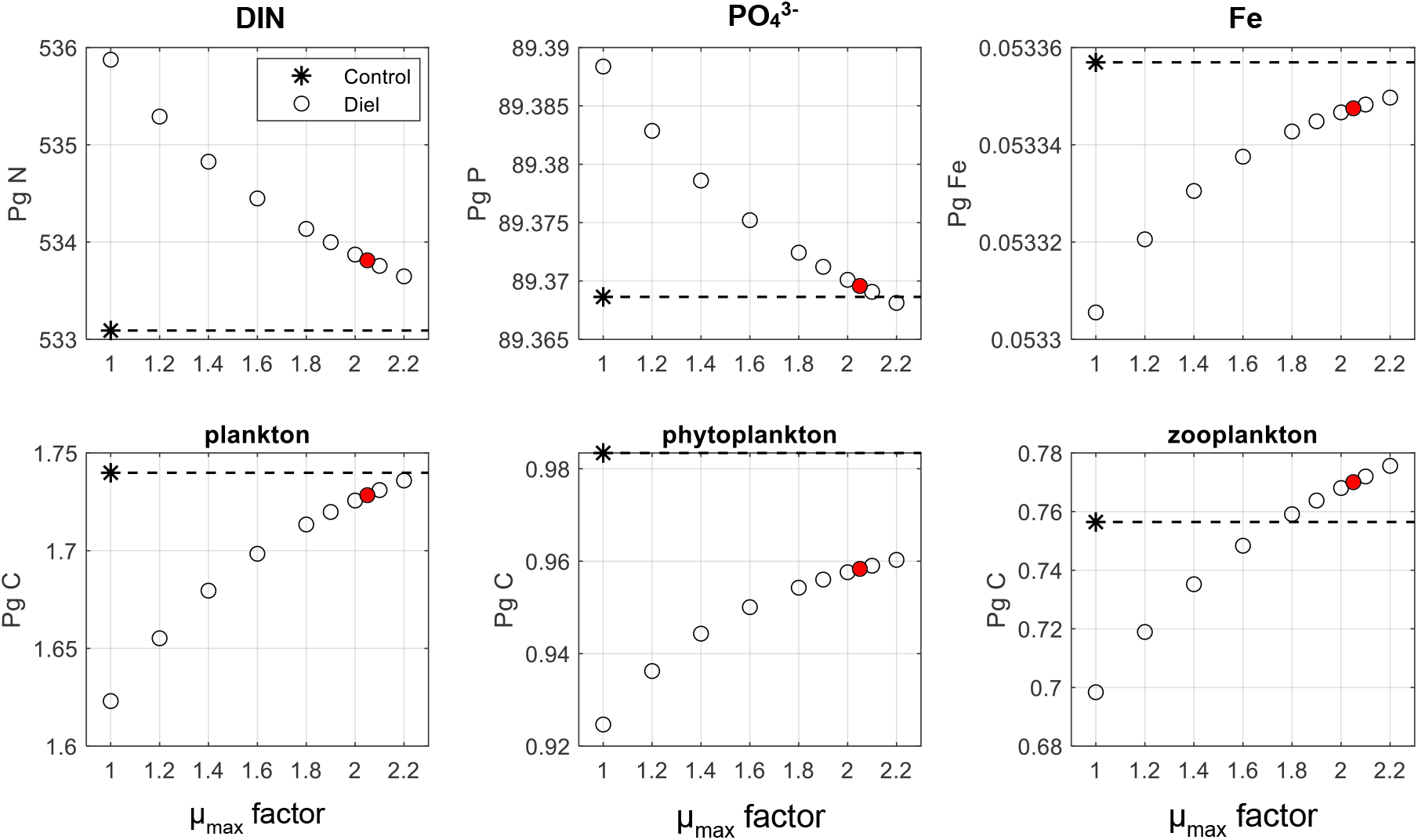
Global ocean inventories of limiting nutrients and phytoplankton biomass of model simulations with only seasonal light cycles (Control) and both seasonal and diel light cycles (Diel). DIN refers to the sum of NO_3_^-^, NO_2_^-^ and NH_4_+. The 10 diel simulations (open circles) differ in phytoplankton growth parameterization; maximal growth rates of all phytoplankton types are multiplied by an μ_max_ factor indicated in x-axis. Note that the diel simulation with μ_max_ factor = 1 is parameterized with same maximal growth rates as the control. The dashed line is an extension of the control, while the red circle highlights the μ_max_ factor = 2.05 where the diel simulation matches the control in global primary production (Fig. 1).

**Figure S2.**
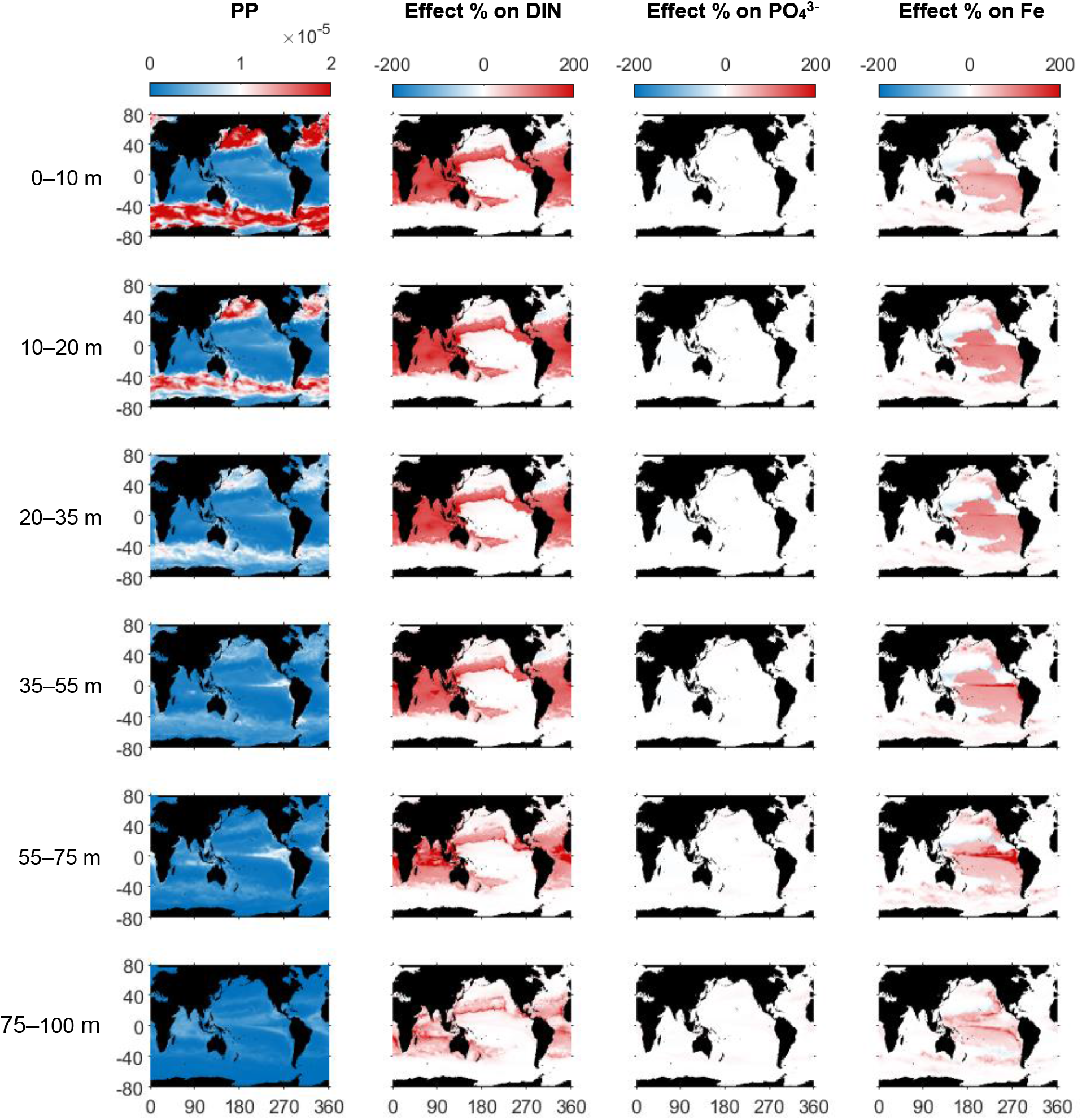
Primary production of the diel simulation, PP (mmol m^-3^ y^-1^) and effect of diel light cycles % compared to the control (Eq. 2) on nutrient concentrations at different model layer depths. DIN refers to the sum of NO_3_^-^, NO_2_^-^ and NH_4_+.

**Figure S3.**
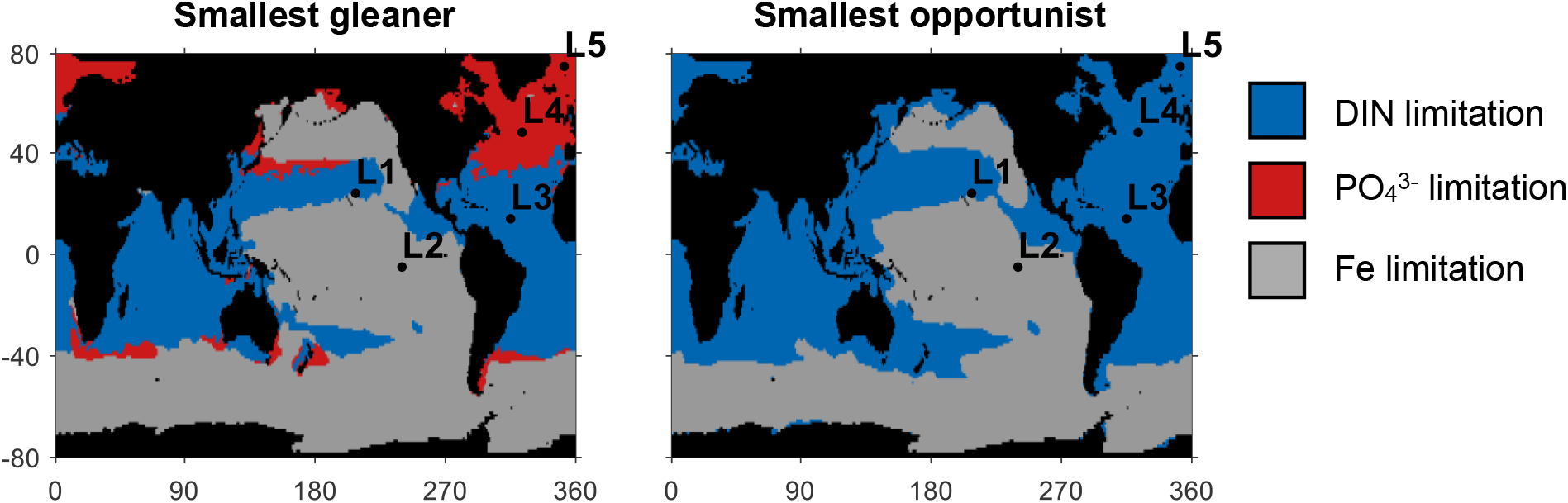
Nutrient limitation in the surface (0 to 55 m) global ocean for the two most abundant phytoplankton types in the model: the smallest gleaner and smallest opportunist. The rest phytoplankton types present similar pattern of nutrient limitation with their smallest group members presented here. Labelled black dots indicate locations (L1 to L5) of which dynamics presented in Fig. 3.

**Figure S4.**
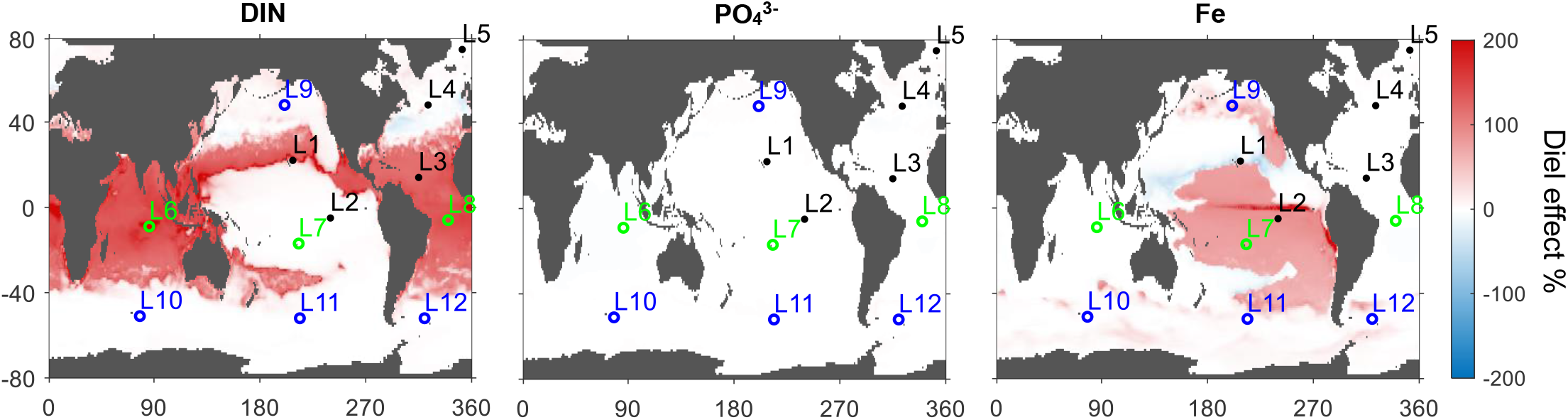
Effect of diel light cycles compared to the control (Eq. 2) on the annually averaged surface (0 – 55 m) concentrations of limiting nutrients (same as lower panel of Fig. 2). DIN refers to the sum of NO_3_^-^, NO_2_^-^ and NH_4_+. Labelled black dots indicate locations (L1 to L5) of which annual dynamics are presented in Fig. 3, while open circles refer to dynamics of additional locations (L6 to L12) presented in Fig. S5: green and blue correspond to tropic and temperate locations, respectively.

**Figure S5.**
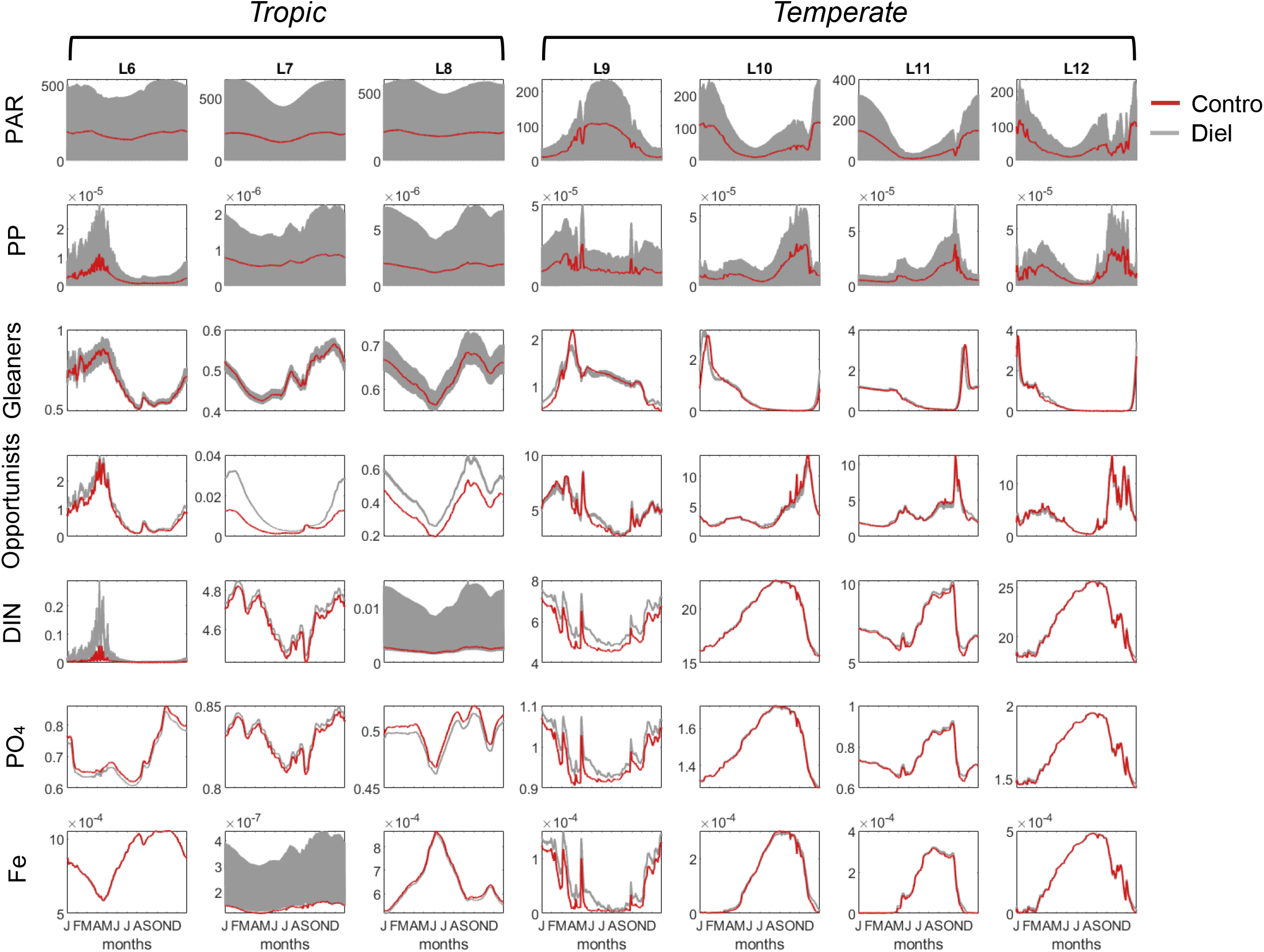
Annual dynamics in the control (red) and diel (grey) simulations of photosynthetically active radiation, PAR (μEinstein m^-2^ s^-1^), primary production, PP (mmol C m^-3^ s^-1^), biomass of phytoplankton gleaners and opportunists (mmol C m^-3^), DIN = NO_3_^-^, NO_2_^-^ and NH_4_+ (mmol N m^-3^), PO_4_^3-^ (mmol P m^-3^) and Fe (mmol Fe m^-3^) in 7 additional locations of the surface ocean (0 to 55 m depth). Locations (L6 to L12) are illustrated in Fig. S4. Note that grey shaded areas indicate diel oscillations.

## Notes

### Competing Interest Statement

The authors have declared no competing interest.

## References

1. Acevedo-Trejos, E., Brandt, G., Merico, A., Smith, S.L., 2013. Biogeographical patterns of phytoplankton community size structure in the oceans. Global Ecology and Biogeography 22, 1060–1070. https://doi.org/10.1111/geb.12071

2. Aumont, O., Maier-Reimer, E., Blain, S., Monfray, P., 2003. An ecosystem model of the global ocean including Fe, Si, P colimitations. Global Biogeochemical Cycles 17. https://doi.org/10.1029/2001GB001745

3. Barton, A.D., Dutkiewicz, S., Flierl, G., Bragg, J., Follows, M.J., 2010. Patterns of Diversity in Marine Phytoplankton. Science 327, 1509–1511. https://doi.org/10.1126/science.1184961

4. Barton, A.D., Pershing, A.J., Litchman, E., Record, N.R., Edwards, K.F., Finkel, Z.V., Kiørboe, T., Ward, B.A., 2013. The biogeography of marine plankton traits. Ecology Letters 16, 522–534. https://doi.org/10.1111/ele.12063

5. Behrenfeld, M.J., Boss, E.S., 2014. Resurrecting the Ecological Underpinnings of Ocean Plankton Blooms. Annual Review of Marine Science 6, 167–194. https://doi.org/10.1146/annurev-marine-052913-021325

6. Boysen, A.K., Carlson, L.T., Durham, B.P., Groussman, R.D., Aylward, F.O., Ribalet, F., Heal, K.R., DeLong, E.F., Armbrust, E.V., Ingalls, A.E., 2020. Diel Oscillations of Particulate Metabolites Reflect Synchronized Microbial Activity in the North Pacific Subtropical Gyre. bioRxiv 2020.05.09.086173. https://doi.org/10.1101/2020.05.09.086173

7. Bracher, A., Vountas, M., Dinter, T., Burrows, J.P., Röttgers, R., Peeken, I., 2009. Quantitative observation of cyanobacteria and diatoms from space using PhytoDOAS on SCIAMACHY data. Biogeosciences 6, 751–764. https://doi.org/10.5194/bg-6-751-2009

8. Brewin, R.J.W., Sathyendranath, S., Hirata, T., Lavender, S.J., Barciela, R.M., Hardman-Mountford, N.J., 2010. A three-component model of phytoplankton size class for the Atlantic Ocean. Ecological Modelling 221, 1472–1483. https://doi.org/10.1016/j.ecolmodel.2010.02.014

9. Brock, T.D., 1981. Calculating solar radiation for ecological studies. Ecological Modelling 14, 1–19. https://doi.org/10.1016/0304-3800(81)90011-9

10. Chai, F., Jiang, M.-S., Chao, Y., Dugdale, R.C., Chavez, F., Barber, R.T., 2007. Modeling responses of diatom productivity and biogenic silica export to iron enrichment in the equatorial Pacific Ocean. Global Biogeochemical Cycles 21. https://doi.org/10.1029/2006GB002804

11. Cochlan, W.P., Harrison, P.J., Denman, K.L., 1991. Diel periodicity of nitrogen uptake by marine phytoplankton in nitrate-rich environments. Limnology and Oceanography 36, 1689–1700. https://doi.org/10.4319/lo.1991.36.8.1689

12. Cohen, S.E., Golden, S.S., 2015. Circadian Rhythms in Cyanobacteria. Microbiol. Mol. Biol. Rev. 79, 373–385. https://doi.org/10.1128/MMBR.00036-15

13. Denny, M., 2017. The fallacy of the average: on the ubiquity, utility and continuing novelty of Jensen’s inequality. Journal of Experimental Biology 220, 139–146. https://doi.org/10.1242/jeb.140368

14. Dutkiewicz, S., Boyd, P.W., Riebesell, U., 2021. Exploring biogeochemical and ecological redundancy in phytoplankton communities in the global ocean. Global Change Biology n/a. https://doi.org/10.1111/gcb.15493

15. Dutkiewicz, S., Cermeno, P., Jahn, O., Follows, M.J., Hickman, A.E., Taniguchi, D.A.A., Ward, B.A., 2020. Dimensions of marine phytoplankton diversity. Biogeosciences 17, 609–634. https://doi.org/10.5194/bg-17-609-2020

16. Dutkiewicz, S., Follows, M.J., Bragg, J.G., 2009. Modeling the coupling of ocean ecology and biogeochemistry. Global Biogeochem. Cycles 23, GB4017. https://doi.org/10.1029/2008GB003405

17. Dutkiewicz, S., Hickman, A.E., Jahn, O., Henson, S., Beaulieu, C., Monier, E., 2019. Ocean colour signature of climate change. Nature Communications 10, 578. https://doi.org/10.1038/s41467-019-08457-x

18. Dutkiewicz, S., Morris, J.J., Follows, M.J., Scott, J., Levitan, O., Dyhrman, S.T., Berman-Frank, I., 2015. Impact of ocean acidification on the structure of future phytoplankton communities. Nature Climate Change 5, 1002–1006. https://doi.org/10.1038/nclimate2722

19. Dutkiewicz, S., Ward, B.A., Scott, J.R., Follows, M.J., 2014. Understanding predicted shifts in diazotroph biogeography using resource competition theory. Biogeosciences 11, 5445–5461. https://doi.org/10.5194/bg-11-5445-2014

20. Edwards, K.F., Thomas, M.K., Klausmeier, C.A., Litchman, E., 2015. Light and growth in marine phytoplankton: allometric, taxonomic, and environmental variation. Limnol. Oceanogr. 60, 540–552. https://doi.org/10.1002/lno.10033

21. Edwards, K.F., Thomas, M.K., Klausmeier, C.A., Litchman, E., 2012. Allometric scaling and taxonomic variation in nutrient utilization traits and maximum growth rate of phytoplankton. Limnol. Oceanogr. 57, 554–566. https://doi.org/10.4319/lo.2012.57.2.0554

22. Finkel, Z.V., Beardall, J., Flynn, K.J., Quigg, A., Rees, T.A.V., Raven, J.A., 2010. Phytoplankton in a changing world: cell size and elemental stoichiometry. J Plankton Res 32, 119–137. https://doi.org/10.1093/plankt/fbp098

23. Flombaum, P., Gallegos, J.L., Gordillo, R.A., Rincón, J., Zabala, L.L., Jiao, N., Karl, D.M., Li, W.K.W., Lomas, M.W., Veneziano, D., Vera, C.S., Vrugt, J.A., Martiny, A.C., 2013. Present and future global distributions of the marine Cyanobacteria Prochlorococcus and Synechococcus. PNAS 110, 9824–9829. https://doi.org/10.1073/pnas.1307701110

24. Follett, C.L., Dutkiewicz, S., Karl, D.M., Inomura, K., Follows, M.J., 2018a. Seasonal resource conditions favor a summertime increase in North Pacific diatom–diazotroph associations. The ISME Journal 12, 1543–1557. https://doi.org/10.1038/s41396-017-0012-x

25. Follett, C.L., White, A.E., Wilson, S.T., Follows, M.J., 2018b. Nitrogen fixation rates diagnosed from diurnal changes in elemental stoichiometry. Limnology and Oceanography 63, 1911–1923. https://doi.org/10.1002/lno.10815

26. Follows, M.J., Dutkiewicz, S., 2011. Modeling Diverse Communities of Marine Microbes. Annual Review of Marine Science 3, 427–451. https://doi.org/10.1146/annurev-marine-120709-142848

27. Follows, M.J., Dutkiewicz, S., Grant, S., Chisholm, S.W., 2007. Emergent Biogeography of Microbial Communities in a Model Ocean. Science 315, 1843–1846. https://doi.org/10.1126/science.1138544

28. Follows, M.J., Dutkiewicz, S., Ward, B., Follett, C., 2018. Theoretical interpretations of subtropical plankton biogeography, in: Gasol, J., Kirchman, D. (Eds.), Microbial Ecology of the Oceans. John Wiley, p. 467.

29. Fuhrman, J.A., Eppley, R.W., Hagström, Å., Azam, F., 1985. Diel variations in bacterioplankton, phytoplankton, and related parameters in the Southern California Bight. Marine Ecology Progress Series 27, 9–20.

30. Geider, R.J., Maclntyre, H.L., Kana, T.M., 1998. A dynamic regulatory model of phytoplanktonic acclimation to light, nutrients, and temperature. Limnology and Oceanography 43, 679–694. https://doi.org/10.4319/lo.1998.43.4.0679

31. Glibert, P.M., Garside, C., 1992. Diel variability in nitrogenous nutrient uptake by phytoplankton in the Chesapeake Bay plume. Journal of Plankton Research 14, 271–288. https://doi.org/10.1093/plankt/14.2.271

32. Grover, J.P., 1990. Resource Competition in a Variable Environment: Phytoplankton Growing According to Monod’s Model. The American Naturalist 136, 771–789.

33. Hansen, P.J., Bjørnsen, P.K., Hansen, B.W., 1997. Zooplankton grazing and growth: Scaling within the 2-2,-μm body size range. Limnol. Oceanogr. 42, 687–704. https://doi.org/10.4319/lo.1997.42.4.0687

34. Hirata, T., Hardman-Mountford, N.J., Brewin, R.J.W., Aiken, J., Barlow, R., Suzuki, K., Isada, T., Howell, E., Hashioka, T., Noguchi-Aita, M., Yamanaka, Y., 2011. Synoptic relationships between surface Chlorophyll-a and diagnostic pigments specific to phytoplankton functional types. Biogeosciences 8, 311–327. https://doi.org/10.5194/bg-8-311-2011

35. Holling, C.S., 1959. Some Characteristics of Simple Types of Predation and Parasitism1. The Canadian Entomologist 91, 385–398. https://doi.org/10.4039/Ent91385-7

36. Holtzendorff, J., Partensky, F., Mella, D., Lennon, J.-F., Hess, W.R., Garczarek, L., 2008. Genome streamlining results in loss of robustness of the circadian clock in the marine cyanobacterium Prochlorococcus marinus PCC 9511. J Biol Rhythms 23, 187–199. https://doi.org/10.1177/0748730408316040

37. Jensen, J.L.W.V., 1906. Sur les fonctions convexes et les inégalités entre les valeurs moyennes. Acta Math. 30, 175–193. https://doi.org/10.1007/BF02418571

38. Johnson, C.H., Zhao, C., Xu, Y., Mori, T., 2017. Timing the day: what makes bacterial clocks tick? Nature Reviews Microbiology 15, 232–242. https://doi.org/10.1038/nrmicro.2016.196

39. Johnson, K.S., Coletti, L.J., Chavez, F.P., 2006. Diel nitrate cycles observed with in situ sensors predict monthly and annual new production. Deep Sea Research Part I: Oceanographic Research Papers 53, 561–573. https://doi.org/10.1016/j.dsr.2005.12.004

40. Johnson, K.S., Coletti, L.J., Jannasch, H.W., Sakamoto, C.M., Swift, D.D., Riser, S.C., 2013. Long-Term Nitrate Measurements in the Ocean Using the in situ Ultraviolet Spectrophotometer: Sensor Integration into the APEX Profiling Float. Journal of Atmospheric and Oceanic Technology 30, 1854–1866. https://doi.org/10.1175/JTECH-D-12-00221.1

41. Kiørboe, T., 1993. Turbulence, Phytoplankton Cell Size, and the Structure of Pelagic Food Webs, in: Blaxter, J.H.S., Southward, A.J. (Eds.), Advances in Marine Biology. Academic Press, pp. 1–72. https://doi.org/10.1016/S0065-2881(08)60129-7

42. Kolody, B.C., McCrow, J.P., Allen, L.Z., Aylward, F.O., Fontanez, K.M., Moustafa, A., Moniruzzaman, M., Chavez, F.P., Scholin, C.A., Allen, E.E., Worden, A.Z., Delong, E.F., Allen, A.E., 2019. Diel transcriptional response of a California Current plankton microbiome to light, low iron, and enduring viral infection. The ISME Journal 13, 2817–2833. https://doi.org/10.1038/s41396-019-0472-2

43. Landolfi, A., Koeve, W., Dietze, H., Kähler, P., Oschlies, A., 2015. A new perspective on environmental controls of marine nitrogen fixation. Geophysical Research Letters 42, 4482–4489. https://doi.org/10.1002/2015GL063756

44. Litchman, E., 2003. Competition and coexistence of phytoplankton under fluctuating light: experiments with two cyanobacteria. Aquatic Microbial Ecology 31, 241–248. https://doi.org/10.3354/ame031241

45. Litchman, E., 1998. Population and community responses of phytoplankton to fluctuating light. Oecologia 117, 247–257. https://doi.org/10.1007/s004420050655

46. Litchman, E., Klausmeier, C.A., 2008. Trait-Based Community Ecology of Phytoplankton. Annual Review of Ecology, Evolution, and Systematics 39, 615–639.

47. Litchman, E., Klausmeier, C.A., 2001. Competition of phytoplankton under fluctuating light. The American Naturalist 157, 170–187.

48. Litchman, E., Klausmeier, C.A., Bossard, P., 2004. Phytoplankton nutrient competition under dynamic light regimes. Limnol. Oceanogr. 49, 1457–1462. https://doi.org/10.4319/lo.2004.49.4_part_2.1457

49. Marañón, E., Cermeño, P., López-Sandoval, D.C., Rodríguez-Ramos, T., Sobrino, C., Huete-Ortega, M., Blanco, J.M., Rodríguez, J., 2013. Unimodal size scaling of phytoplankton growth and the size dependence of nutrient uptake and use. Ecol Lett 16, 371–379. https://doi.org/10.1111/ele.12052

50. Marshall, J., Adcroft, A., Hill, C., Perelman, L., Heisey, C., 1997. A finite-volume, incompressible Navier Stokes model for studies of the ocean on parallel computers. Journal of Geophysical Research: Oceans 102, 5753–5766. https://doi.org/10.1029/96JC02775

51. Martiny, A.C., Tai, A.P.K., Veneziano, D., Primeau, F., Chisholm, S.W., 2009. Taxonomic resolution, ecotypes and the biogeography of Prochlorococcus. Environmental Microbiology 11, 823–832. https://doi.org/10.1111/j.1462-2920.2008.01803.x

52. Monod, J., 1949. The Growth of Bacterial Cultures. Annual Review of Microbiology 3, 371–394. https://doi.org/10.1146/annurev.mi.03.100149.002103

53. Needoba, J.A., Harrison, P.J., 2004. Influence of Low Light and a Light: Dark Cycle on No3– Uptake, Intracellular No3–, and Nitrogen Isotope Fractionation by Marine Phytoplankton1. Journal of Phycology 40, 505–516. https://doi.org/10.1111/j.1529-8817.2004.03171.x

54. Prowe, A.E.F., Pahlow, M., Dutkiewicz, S., Follows, M., Oschlies, A., 2012. Top-down control of marine phytoplankton diversity in a global ecosystem model. Progress in Oceanography 101, 1–13. https://doi.org/10.1016/j.pocean.2011.11.016

55. Ptacnik, R., Solimini, A.G., Andersen, T., Tamminen, T., Brettum, P., Lepistö, L., Willén, E., Rekolainen, S., 2008. Diversity predicts stability and resource use efficiency in natural phytoplankton communities. PNAS 105, 5134–5138. https://doi.org/10.1073/pnas.0708328105

56. Raven, J.A., 1994. Why are there no picoplanktonic O2 evolvers with volumes less than 10-19 m3? J Plankton Res 16, 565–580. https://doi.org/10.1093/plankt/16.5.565

57. Sakamoto, C.M., Johnson, K.S., Coletti, L.J., Maurer, T.L., Massion, G., Pennington, J.T., Plant, J.N., Jannasch, H.W., Chavez, F.P., 2017. HOURLY IN SITU NITRATE ON A COASTAL MOORING: A 15-Year Record and Insights into New Production. Oceanography 30, 114–127.

58. Sommer, U., Adrian, R., Domis, L.D.S., Elser, J.J., Gaedke, U., Ibelings, B., Jeppesen, E., Lürling, M., Molinero, J.C., Mooij, W.M., Donk, E. van, Winder, M., 2012. Beyond the Plankton Ecology Group (PEG) Model: Mechanisms Driving Plankton Succession. Annual Review of Ecology, Evolution, and Systematics 43, 429–448. https://doi.org/10.1146/annurev-ecolsys-110411-160251

59. Sommer, U., Charalampous, E., Genitsaris, S., Moustaka-Gouni, M., 2017. Benefits, costs and taxonomic distribution of marine phytoplankton body size. J Plankton Res 39, 494–508. https://doi.org/10.1093/plankt/fbw071

60. Stramska, M., Dickey, T.D., 1992. Variability of bio-optical properties of the upper ocean associated with diel cycles in phytoplankton population. Journal of Geophysical Research: Oceans 97, 17873–17887. https://doi.org/10.1029/92JC01570

61. Thomas, D.N., 2003. Iron Limitation in the Southern Ocean. Science 302, 565–566. https://doi.org/10.1126/science.302.5645.565c

62. Tilman, D., 1982. Resource competition and community structure. Princeton University Press.

63. Tréguer, P., Bowler, C., Moriceau, B., Dutkiewicz, S., Gehlen, M., Aumont, O., Bittner, L., Dugdale, R., Finkel, Z., Iudicone, D., Jahn, O., Guidi, L., Lasbleiz, M., Leblanc, K., Levy, M., Pondaven, P., 2018. Influence of diatom diversity on the ocean biological carbon pump. Nature Geoscience 11, 27–37. https://doi.org/10.1038/s41561-017-0028-x

64. Tsakalakis, I., Follows, M.J., Dutkiewicz, S., Follett, C.L., Vallino, J.J., 2021. Global ocean model results – effects of diel light cycles. Zenodo. https://doi.org/10.5281/zenodo.4772367

65. Tsakalakis, I., Pahlow, M., Oschlies, A., Blasius, B., Ryabov, A.B., 2018. Diel light cycle as a key factor for modelling phytoplankton biogeography and diversity. Ecological Modelling 384, 241–248. https://doi.org/10.1016/j.ecolmodel.2018.06.022

66. Vallina, S.M., Follows, M.J., Dutkiewicz, S., Montoya, J.M., Cermeno, P., Loreau, M., 2014. Global relationship between phytoplankton diversity and productivity in the ocean. Nature Communications 5, 4299. https://doi.org/10.1038/ncomms5299

67. Vallino, J.J., Tsakalakis, I., 2020. Phytoplankton Temporal Strategies Increase Entropy Production in a Marine Food Web Model. Entropy 22, 1249. https://doi.org/10.3390/e22111249

68. Ward, B.A., Dutkiewicz, S., Jahn, O., Follows, M.J., 2012. A size-structured food-web model for the global ocean. Limnol. Oceanogr. 57, 1877–1891. https://doi.org/10.4319/lo.2012.57.6.1877

69. Ward, B.A., Follows, M.J., 2016. Marine mixotrophy increases trophic transfer efficiency, mean organism size, and vertical carbon flux. PNAS 113, 2958–2963. https://doi.org/10.1073/pnas.1517118113

70. Winder, M., Sommer, U., 2012. Phytoplankton response to a changing climate. Hydrobiologia 698, 5–16. https://doi.org/10.1007/s10750-012-1149-2

71. Wunsch, C., Heimbach, P., 2007. Practical global oceanic state estimation. Physica D: Nonlinear Phenomena, Data Assimilation 230, 197–208. https://doi.org/10.1016/j.physd.2006.09.040

72. Yingling, N., Kelly, T.B., Shropshire, T.A., Landry, M.R., Selph, K.E., Knapp, A.N., Kranz, S.A., Stukel, M.R., 2021. Taxon-specific phytoplankton growth, nutrient utilization, and light limitation in the oligotrophic Gulf of Mexico. bioRxiv 2021.03.01.433426. https://doi.org/10.1101/2021.03.01.433426

73. Zhang, J.-Z., Wanninkhof, R., Lee, K., 2001. Enhanced new production observed from the diurnal cycle of nitrate in an oligotrophic anticyclonic eddy. Geophysical Research Letters 28, 1579–1582. https://doi.org/10.1029/2000GL012065

